# Molecular basis of EMRE-dependence of the human mitochondrial calcium uniporter

**DOI:** 10.1101/637918

**Authors:** Melissa J.S. MacEwen, Andrew L. Markhard, Mert Bozbeyoglu, Forrest Bradford, Olga Goldberger, Vamsi K. Mootha, Yasemin Sancak

**Affiliations:** Department of Pharmacology, University of Washington, Seattle, WA, USA; Howard Hughes Medical Institute and Department of Molecular Biology, Massachusetts General Hospital, Boston, MA, USA; Broad Institute, Cambridge, MA, USA; Department of Systems Biology, Harvard Medical School, Boston, MA, USA

## Abstract

The mitochondrial uniporter is calcium-activated calcium channel complex critical for cellular signaling and bioenergetics. MCU, the pore-forming subunit of the uniporter, contains two transmembrane domains and is found in all major eukaryotic taxa. In amoeba and fungi, MCU homologs are sufficient to form a functional calcium channel, whereas human MCU exhibits a strict requirement for the metazoan-specific, single-pass transmembrane protein EMRE for conductance. Here, we exploit this evolutionary divergence to decipher the molecular basis of the human MCU’s dependence on EMRE. By systematically generating chimeric proteins that consist of EMRE-independent *D. discoideum* MCU (DdMCU) and *H. sapiens* MCU (HsMCU), we converged on a stretch of 10 amino acids in DdMCU that can be transplanted to HsMCU to render it EMRE-dependent. We call this region in human MCU the EMRE-dependence domain (EDD). Crosslinking experiments show that HsEMRE directly interacts with MCU at both of its transmembrane domains as well as the EDD. Based on previously published structures of fungal MCU homologs, the EDD segment is located distal to the calcium pore’s selectivity filter and appears flexible. We propose that EMRE stabilizes EDD of MCU, permitting both channel opening and calcium conductance

## INTRODUCTION

Mitochondria play central roles in diverse cellular processes, including metabolism, signaling and cell death. Calcium (Ca^2+^) signaling is critical in coordination of cellular needs with mitochondrial outputs by regulating the activity of the tricarboxylic acid (TCA) cycle, activity of mitochondrial metabolite carriers and triggering the mitochondrial permeability transition pore (Del Arco, Contreras, Pardo, & Satrustegui, 2016; Denton, 2009; Giorgio, Guo, Bassot, Petronilli, & Bernardi, 2018). This coordination is partially mediated by entry of Ca^2+^ into the mitochondrial matrix from the cytosol during a Ca^2+^ signaling event. Perturbation of mitochondrial Ca^2+^ uptake is associated with a plethora of cellular and systemic pathologies, ranging from abnormal mitochondrial movement and shape, to immune dysfunction, cell cycle progression and neuromuscular disease (Koval et al., 2019; Logan et al., 2014; Mammucari et al., 2015; Mammucari et al., 2018; Paupe & Prudent, 2018; Prudent et al., 2016; Zhao et al., 2019).

The mitochondrial Ca^2+^ uniporter complex, a multi-subunit protein assembly that resides in the inner mitochondrial membrane (IMM), is responsible for bulk entry of Ca^2+^ ions into the mitochondrial matrix ((Carafoli & Lehninger, 1971; Deluca & Engstrom, 1961; Kirichok, Krapivinsky, & Clapham, 2004; Vasington & Murphy, 1962). A ~35kDa protein termed MCU (mitochondrial calcium uniporter) is the defining component of the uniporter complex and serves as its pore (Baughman et al., 2011; Chaudhuri, Sancak, Mootha, & Clapham, 2013; De Stefani, Raffaello, Teardo, Szabo, & Rizzuto, 2011; Kovacs-Bogdan et al., 2014). MCU is a transmembrane protein with two membrane-spanning helices (TM1 and TM2), a short linker region facing the intermembrane with a “DIME” motif, a large amino terminal domain that assumes a β-grasp fold (S. K. Lee et al., 2016; Y. Lee et al., 2015) and a carboxyl terminal region that is mostly helical (Baradaran, Wang, Siliciano, & Long, 2018; Fan et al., 2018; Nguyen et al., 2018; Oxenoid et al., 2016; Yoo et al., 2018). Functional and structural studies showed that TM2 forms the Ca^2+^-conducting pore of the channel, whereas the N-terminal domain is mostly dispensable for Ca^2+^ conductance and is likely to play a regulatory role. In animals, MCU nucleates other proteins (MCUb, MICU1, MICU2, MICU3, EMRE) that regulate different aspects of uniporter function. MICU1, MICU2 and MICU3 are EF-hand containing Ca^2+^-binding proteins that localize to the intermembrane space (IMS) of the mitochondria. None of the MICU homologs is necessary for Ca^2+^ conductance by the uniporter, but rather, they play crucial roles in setting the threshold for Ca^2+^ uptake (Csordas et al., 2013; de la Fuente, Matesanz-Isabel, Fonteriz, Montero, & Alvarez, 2014; Kamer et al., 2018; Kevin Foskett & Madesh, 2014; Liu et al., 2016; Mallilankaraman et al., 2012; Patron et al., 2014; Perocchi et al., 2010; Plovanich et al., 2013). MCUb is a paralog of MCU and is thought to be a negative regulator of MCU due to its inability to form a functional Ca2+ channel (Raffaello et al., 2013).

EMRE is a single-pass transmembrane protein that was the last component of the uniporter to be identified, in part because of its curious evolutionary distribution(Sancak et al., 2013). MCU and MICU1 homologs tend to be found in all major eukaryotic taxa, with lineage specific losses(Bick, Calvo, & Mootha, 2012). S. *cerevisiae*, for example, has completely lost both MICU1 and MCU, and in fact, this evolutionary diversity formed the basis for the initial discovery of MICU1 (Perocchi et al., 2010). Following the initial molecular identification of the uniporter machinery, efforts to functionally reconstitute uniporter activity in yeast mitochondria using human MCU alone failed, for reasons that were not clear. This led to the search for additional missing components of the uniporter complex, leading to the identification of EMRE, which is present only in metazoans (Sancak et al., 2013). In these species, EMRE fulfills two important functions. First, the C-terminal domain of EMRE is crucial for MCU-MICU1 interaction (Sancak et al., 2013; M. F. Tsai et al., 2016). Second, EMRE is strictly required for mitochondrial Ca^2+^ uptake (Kovacs-Bogdan et al., 2014; Sancak et al., 2013; M. F. Tsai et al., 2016). Hence, in metazoans, MCU and EMRE are both necessary for uniporter pore activity. However, the mechanism by which EMRE enables Ca^2+^ conductance by the metazoan MCU has remained unknown.

Here, we exploited the evolutionary divergence of EMRE to understand its role in the uniporter complex. We previously showed that in slime mold *Dictyostelium discoideum*, there are no EMRE homologs and *Dictyostelium discoideum* MCU (DdMCU) forms a functional uniporter by itself (Kovacs-Bogdan et al., 2014). We reasoned that sequence elements that confer EMRE-independent activity to DdMCU could be swapped from DdMCU to HsMCU to render it EMRE-independent. To this end, we systematically generated HsMCU-DdMCU chimeric proteins and tested their ability to conduct Ca^2+^ in the absence of EMRE. These efforts led to the identification of a 10 amino acid-long region in HsMCU that determines its EMRE-dependence. We call this region EMRE Dependence Domain (EDD). Using copper-mediated cysteine crosslinking experiments, we show that EMRE interacts with both transmembrane domains of MCU (TM1 and TM2) as well as its EDD. Interestingly, EDD, which is C-terminal to the pore-forming TM2, appears flexible in published high-resolution MCU structures. Our data suggest that EMRE stabilizes this region through direct binding, which leads to the formation of a functional channel in metazoans.

## RESULTS

### Carboxyl-terminal domain of EMRE faces the intermembrane space and mediates MICU1-EMRE interaction

EMRE is a small transmembrane protein that resides in the inner mitochondrial membrane and has been shown to have two distinct functions in the uniporter. First, EMRE facilitates the interaction of MCU with MICU1. Second, it is required for Ca^2+^ conductance through human MCU. It was essential to clarify EMRE’s membrane topology to understand the mechanisms of these two functions. To this end, we first generated EMRE KO cell lines that stably express EMRE protein tagged with FLAG at its carboxyl terminus (C-terminus) (EMRE-FLAG). When expressed at endogenous levels, EMRE-FLAG rescued the mitochondrial Ca^2+^ uptake defect observed in EMRE KO cells to the same extend as untagged EMRE protein (Fig1A) in a permeabilized cell mitochondrial Ca^2+^uptake assay. Furthermore, EMRE-FLAG immunoprecipitated endogenous MCU and MICU1 (Figure 1B), showing that the C-terminal tag did not perturb EMRE’s function or interaction with other uniporter proteins.

**Figure 1:**
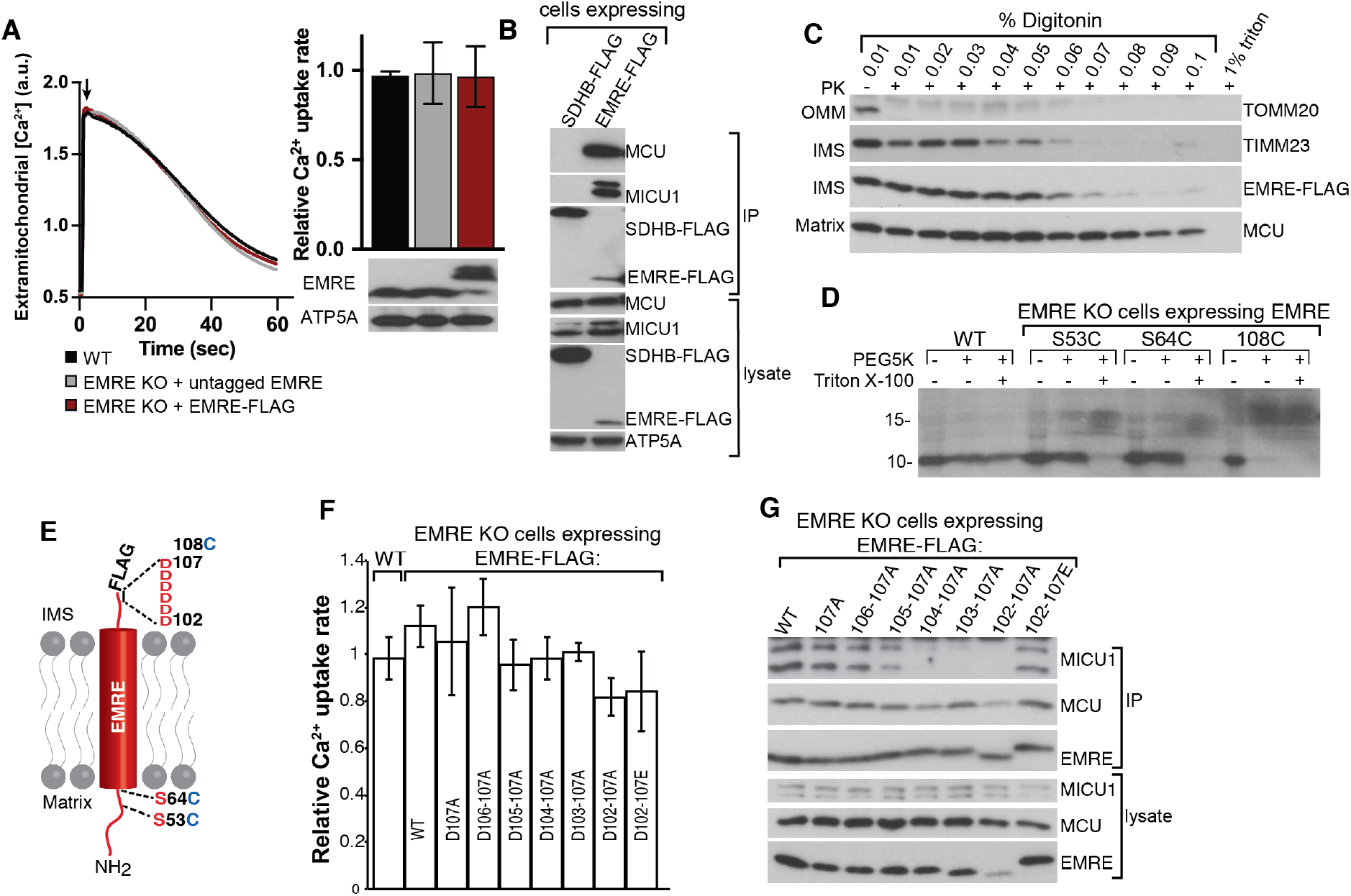
EMRE CAD faces the IMS and mediates EMRE-MICU1 interaction. (A) Tagging EMRE with FLAG epitope tag at its C terminus does not impair its function. HEK293T cells expressing indicated proteins were permeabilized and mitochondiral Ca^2+^ uptake was measured by monitoring extramitochondrial Ca^2+^ clearance. Bar graph shows quantification of Ca^2+^ uptake rates and western blot shows EMRE expression. ATP5A serves as loading control. (B) C-terminal FLAG tag does not impair EMRE-MCU and EMRE-MICU1 interactions. EMRE FLAG and control SDHB-FLAG were immunoprecipitated and immunoprecipitates were blotted for MCU and MICU1. ATP5A serves as loading control. (C) Proteinase K treatment of isolated mitochondria in the presence of increased detergent concentration. EMRE-FLAG is degraded by PK at the same detergent concentration as TIMM23, an inner mitochondrial membrane protein. (D) Mitochondria were isolated from WT or EMRE KO cells that stably express the indicated proteins. Mitoplasts (mitochondria without outer membranes) were prepared and treated with PEG5K-maleimide. A 5kDa mass addition to EMRE protein was detected by western blotting. (E) Schematic shows EMRE membrane topology and the position of the amino acids that were mutated to cysteines for PEGylation experiments shown in (D). EMRE aa 64-85 were predicted to form its transmembrane domain using TMHMM (Sonnhammer, von Heijne, & Krogh, 1998).

To determine the membrane topology of EMRE, we first used a proteinase accessibility assay. Mitochondria isolated from EMRE-FLAG expressing EMRE KO cells were incubated with proteinase K in the presence of increasing concentrations of digitonin, and degradation of EMRE-FLAG was monitored using western blotting. The FLAG tag disappeared at the same digitonin concentration as IMS protein TIMM23, suggesting that the C-terminus of EMRE faces the IMS (Figure 1C). Next, we confirmed N-in C-out topology of EMRE by using an orthogonal approach that utilizes the addition of a 5 kiloDalton (kDa) mass to cysteine residues using polyethylene glycol (PEG)-maleimide (PEG5K). In this assay, cysteine residues that are in the matrix are shielded from membrane impermeable PEG5K. Wild type EMRE does not contain any cysteines, so we mutated S53 or S64 – amino acids that are N-terminal to the predicted transmembrane domain (aa 65-84) – to cysteine (S53C or S64C). We also added a cysteine residue at EMRE’s C-terminus after the last amino acid (EMRE 108C). Expression of WT, S53C, S64C or 108C EMRE in EMRE KO cells rescued mitochondrial Ca^2+^ uptake defect of these cells **(Supplementary Figure 1)**, suggesting that these mutations do not perturb protein function and topology. Mitoplasts (mitochondria without an outer membrane) prepared from EMRE KO cells that express wild type, S53C, S63C or 108C EMRE were treated with PEG5K-maleimide. A ~5kDa shift on molecular weight of EMRE was detected only with the EMRE108C protein, suggesting that 53C and 64C are in the matrix. When PEG5K was added in the presence of a small amount of detergent to disrupt the inner membrane, all three cysteine-containing proteins were PEGylated, showing that the lack of modification of 53C and 64C was not due to their inaccessibility to PEG5K-maleimide in the complex (Figure 1D). These findings are consistent with previous results and confirm that EMRE’s N terminus faces the matrix and its C-terminus acidic domain (CAD) faces the IMM (Figure 1E) (M. F. Tsai et al., 2016; Yamamoto et al., 2016).

CAD has a high percentage of negatively charged aspartic acid (D) and glutamic acid (E) (10/22 amino acids, ~45%). Notably, the presence of five or more D or E at the end of the protein is conserved across species and is a defining feature of EMRE (Sancak et al). CAD has been shown to be important for the interaction of EMRE with MICU1. We asked whether EMRE-MICU1 interaction is mainly mediated by the negative charges in this region and whether CAD also plays a role in Ca^2+^conductance. To test these, we mutated the six Ds (D102 to D107) to alanine (A) and expressed these mutant proteins in EMRE KO cells as FLAG-tagged proteins. Loss of negative charge in this region did not perturb mitochondrial Ca^2+^ uptake (Figure 1F) (M. F. Tsai et al., 2016; Yamamoto et al., 2016). However, we did see a gradual decrease in the amount of MICU1 that immunoprecipitated with EMRE as the number of alanines increased in this region. EMRE-MICU1 interaction was restored when Ds were mutated to similarly charged Es (Figure 1G). These results show that the negative charge of CAD is dispensable for Ca^2+^ conductance, but is critical for EMRE-MICU1 interaction.

### Identification of EMRE-dependence domain (EDD) of HsMCU

EMRE is a metazoan-specific protein, and its loss leads to a complete loss of mitochondrial Ca^2+^ uptake, but the molecular basis for this requirement is not known. We previously showed that slime mold *Dictyostelium discoideum* does not have an EMRE homolog and that D. *discoideum* MCU (DdMCU) forms a functional Ca^2+^ channel by itself (Kovacs-Bogdan et al., 2014). In contrast, to be able to conduct Ca^2+^, human MCU (HsMCU) requires co-expression of EMRE. EMRE-dependence of MCU does not appear to be related to proper MCU folding or mitochondrial localization, as MCU forms higher order oligomers with correct membrane topology in the absence of EMRE (Kovacs-Bogdan et al., 2014). We hypothesized that EMRE plays an important role in Ca^2+^ permeation of the human uniporter, and exploited the evolutionarily divergence of EMRE-dependence of DdMCU and HsMCU to understand the molecular details of EMRE function. First, we compared the predicted secondary structures of DdMCU and HsMCU (Figure 2A). The predicted secondary structures of the two MCU proteins were most divergent at their N-termini. To test whether the DdMCU N-terminus domain would be sufficient to confer EMRE independence to HsMCU, we generated a chimeric protein, chimera 1, as shown in Figure 2B. We expressed chimera 1 in MCU KO HEK293T cells to determine whether it would form a functional protein, and in EMRE KO HEK293T cells to determine whether it would function independently of EMRE in our permeabilized cell mitochondrial Ca^2+^ uptake assay. Chimera 1 formed a functional channel in MCU KO cells, however, its activity was still dependent on the presence of EMRE (Figure 2B).

**Figure 2:**
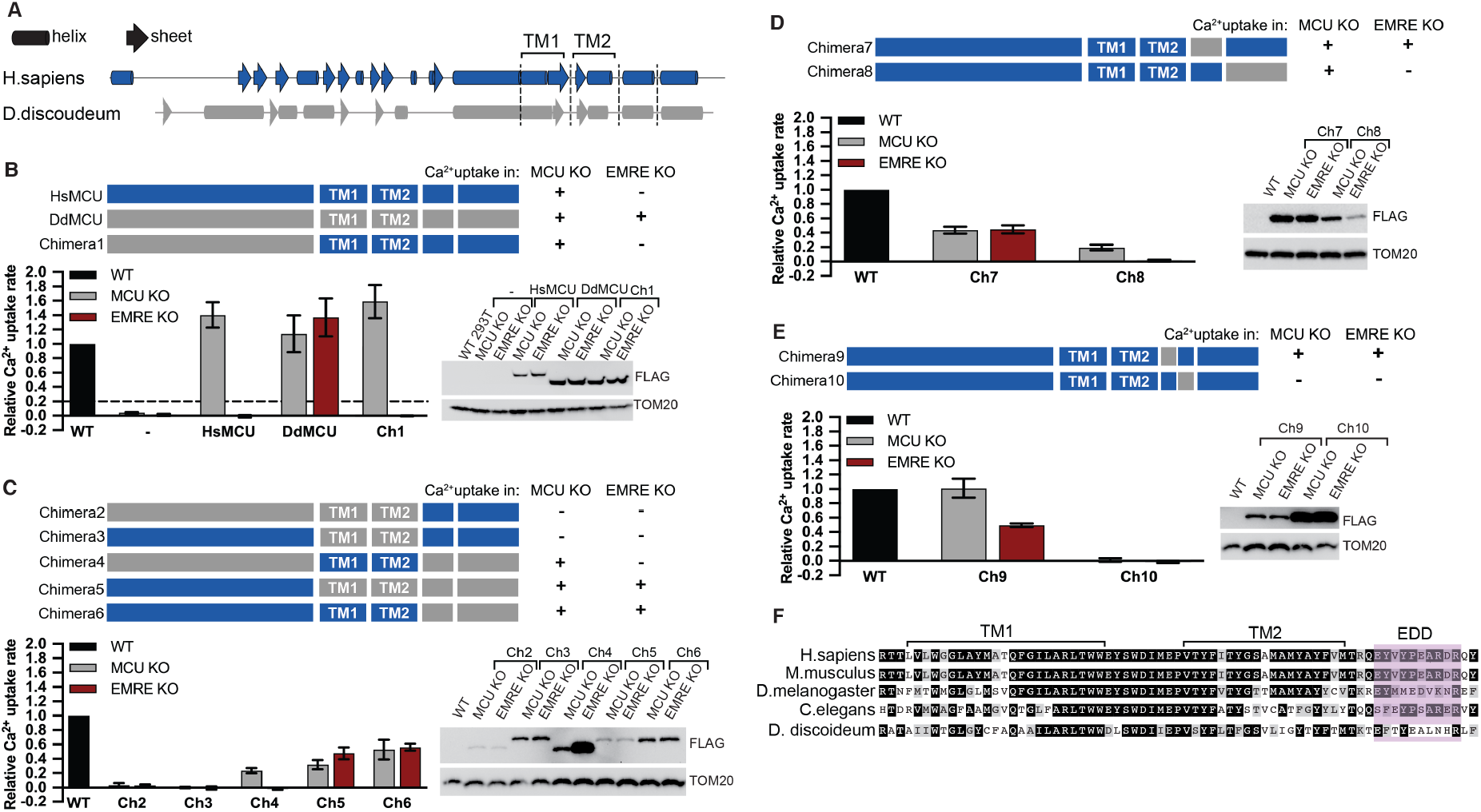
EMRE-Dependence Domain (EDD) of HsMCU is a 10-amino acid long region located C-terminal to TM2. (A) Schematic shows helices and sheets of HsMCU and DdMCU as predicted by PSIPRED. Two transmembrane domains are labeled. (B) Schematic summarizes the domain structure of the chimeric proteins. FLAG-tagged chimeras were stably expressed in MCU KO and EMRE KO HEK293T cells. Mitochondrial Ca^2+^ uptake rates in control WT and chimera expressing cells were measured and normalized to those of WT cells. Error bars report -/+S.D, n=3 or 4. Expression of chimeras was detected by western blotting using anti-FLAG antibody. TOM20 serves as loading control. (C)-(E) Schematic summarizing the domain structure of the chimeric proteins. FLAG-tagged chimeras were stably expressed in MCU KO and EMRE KO HEK293T cells. Mitochondrial Ca^2+^ uptake rates in control WT and chimera expressing cells were measured and normalized to those of WT cells. Expression of chimeras was detected by western blotting using anti-FLAG antibody. TOM20 serves as loading control. (F) Alignment of MCU protein from indicated species was done using CLUSTALW and amino acids were color coded using Boxshade. Black boxes show identical amino acids, gray boxes show similar amino acids. TM1, TM2 and EDD are indicated.

These results suggested that the two transmembrane domains or the C-terminus domain of HsMCU might be critical for its EMRE-dependence. To test this, we generated chimeras 2-6 and determined their functionality and EMRE-dependence. Chimeras 2 and 3 did not form functional Ca^2+^ channels (Figure 2C). Chimera 4 showed reduced, but EMRE-dependent uniporter activity (Figure 2C), suggesting that the TM domains of DdMCU alone are not sufficient for EMRE-independent Ca^2+^ uptake. Chimeras 5 and 6, on the other hand, showed EMRE-independent Ca^2+^ uptake (Figure 2C). Both of these chimeras contain the two predicted C-terminal helices from DdMCU. To determine whether one of these predicted helices might be the critical domain for EMRE-independent Ca^2+^uptake, we generated chimeras 7 and 8. Surprisingly, chimera 7 was functional to the same extent both in MCU KO and EMRE KO cells. In addition, chimera 8, despite its poor expression, supported mitochondrial Ca^2+^ uptake, and it’s activity was EMRE-dependent (Figure 2D). The helical region that defines chimera 7 is composed of 23 amino acids. We further divided this region into two halves at conserved amino acids to generate chimeras 9 and 10 (Figure 2E, **Supplementary Figure 2**). Chimera 10 did not support mitochondrial Ca^2+^ uptake, but chimera 9 formed an EMRE-independent Ca^2+^ channel (Figure 2E). The chimera 9 region originating from DdMCU is 10 amino acids long and is located directly after the pore forming TM (TM2) of HsMCU. We term this 10 amino acid region of MCU (aa288-aa297) the EMRE dependence domain (EDD) (Figure 2F).

During our analysis, we noticed that the expression levels of the chimeric proteins varied (Figures 2B-2D, Supplementary Figure 3A). However, we did not observe a correlation between the expression level of a particular chimera and the rate of mitochondrial Ca^2+^ uptake in cells that express the chimeric proteins (for example, compare the chimera 4 and 5 in Figure 2C). In these experiments, chimeric proteins were stably expressed in cells using lentivirus-mediated integration of the corresponding cDNAs into the genome. To eliminate the possibility that low protein expression was due to low virus titer, we picked chimera 5, a chimera that expressed poorly but formed functional channels, and re-infected cells with virus. We observed a slight increase in protein expression and a concomitant increase in Ca^2+^ uptake rates. However, chimera 5 still expressed at lower levels compared to HsMCU **(Supplementary Figure 3B)**. This suggests that chimeric proteins may be inherently unstable. Consequently, we cannot determine whether low Ca^2+^ uptake rates of chimeras compared to HsMCU are due to their channel properties or their expression levels. Nevertheless, expression of a particular chimeric protein in MCU KO and EMRE KO cells were mostly comparable, allowing us to determine their EMRE-dependence. Expression of HsMCU-DdMCU chimeras did not alter mitochondrial membrane potential; suggesting that lack of mitochondrial Ca^2+^ uptake after expression of some chimeras was not secondary to perturbed mitochondrial health **(Supplementary Figure 4)**. Example mitochondrial Ca^2+^ uptake data for HsMCU, DdMCU and chimeras in MCU KO and EMRE KO cells are shown in **Supplementary Figure 5** and sequences of the chimeric proteins used are provided (Supplementary Materials).

### MCU TM1, TM2 and EDD interact with EMRE

Our functional experiments highlighted the importance of MCU TM domains and EDD for the EMRE-dependence of human MCU. To determine if these domains are also important for EMRE-MCU interaction, we performed immunoprecipitation experiments using chimeras that formed functional channels and showed good protein expression (chimeras 1, 5, 6, 7 and 9). To further normalize protein expression levels, we transiently expressed control HsMCU, DdMCU or chimeras, together with untagged EMRE in MCU KO cells. We treated these cells with amine-reactive crosslinker dithiobis(succinimidyl propionate) (DSP) DSP before cell lysis to stabilize protein-protein interactions. We then immunoprecipitated MCU-FLAG. Chimera 5 did not interact with EMRE; chimeras 6 and 7 showed reduced EMRE interaction; chimeras 1 and 9 showed wild type levels of EMRE-MCU interaction (Figures 3A and 3B). Consistent with stabilization of EMRE protein when bound to MCU, chimeras that showed better EMRE interaction also had more EMRE protein in the lysate (Figure 3A). These experiments suggested that MCU TM1, TM2 and the helical region that includes the EDD are important for EMRE-MCU binding. To determine if EMRE directly interacts with HsMCU in TM1, TM2 and EDD, we performed cysteine-crosslinking experiments.

**Figure 3:**
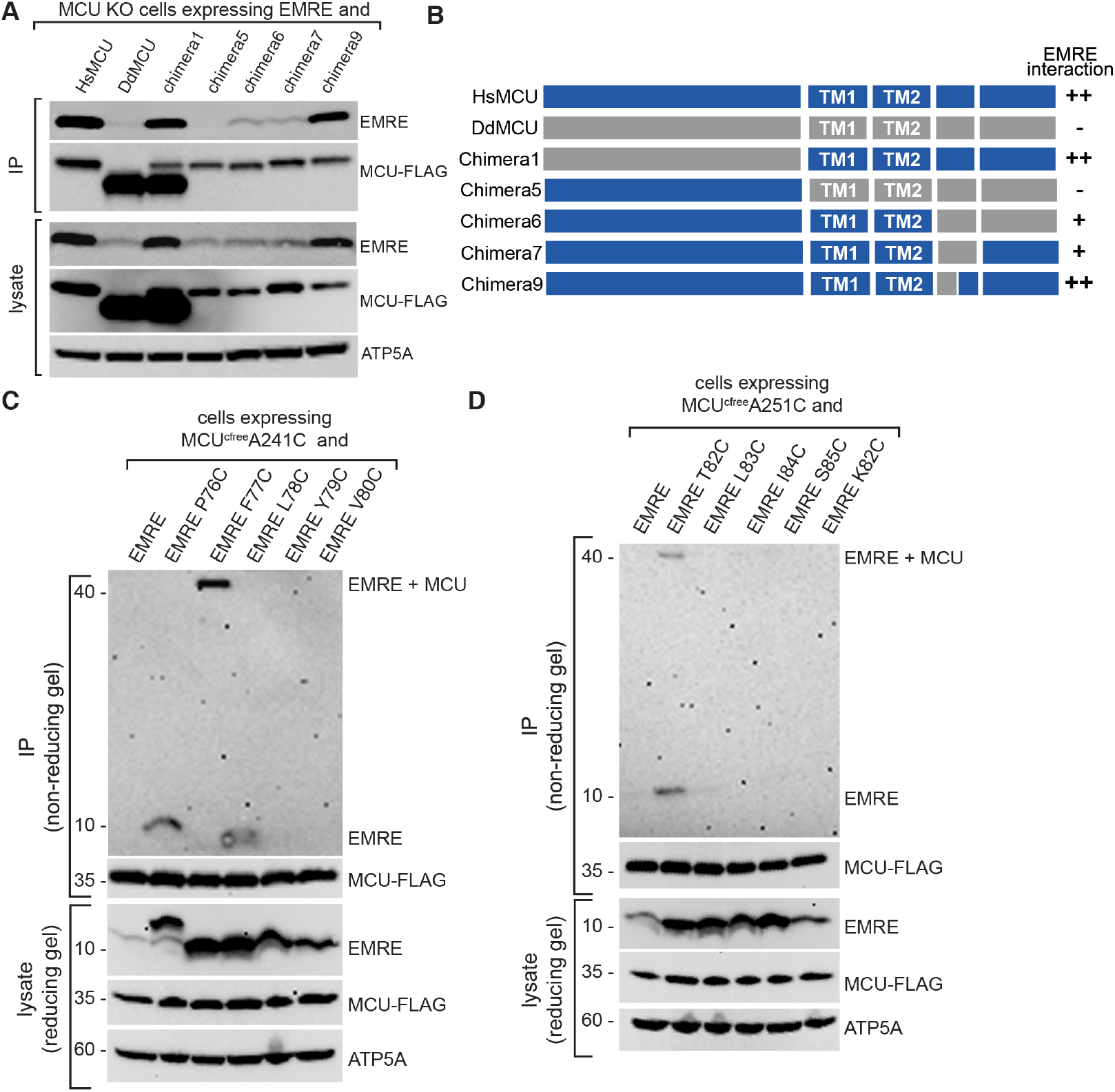
EMRE directly interacts with TM1 of MCU. (A) MCU TM and C-terminal helices are required for EMRE-MCU interaction. Untagged EMRE and indicated FLAG-tagged MCU proteins were co-expressed in MCU KO HEK293T cells by transient transfection, FLAG-tagged proteins were immunoprecipitated and immunoprecipitates were analyzed for the presence of EMRE by western blotting. ATP5A serves as loading control. EMRE-MCU interaction was evident both in immunoprecipitates and in lysates through stabilization of EMRE. (B) Schematic summarizes EMRE-chimera binding data and highlights the importance of MCU TM and C-terminal helices for EMRE-MCU interaction. (C)-(D) EMRE-HsMCU cysteine crosslinking experiments show direct binding of MCU TM1 residues A241 and A251 to EMRE F77 and T82, respectively. HsMCU that contains only one cysteine at amino acid 241 (C) or 251 (D) were stably co-expressed with indicated EMRE proteins. WT EMRE does not contain any cysteines and served as a control. Mitochondria were isolated from cells, and cysteine-cysteine crosslinking was induced using copper phenanthroline. MCU-FLAG was immunoprecipitated and the presence of a ~40 kDa crosslinked EMRE-MCU band was detected under non-reducing conditions by western blotting using EMRE antibody. Lysates were prepared in parallel under reducing conditions and were blotted with indicated proteins. ATP5A serves as a loading control. Numbers indicate the locations of molecular weight standards.

**Figure 4:**
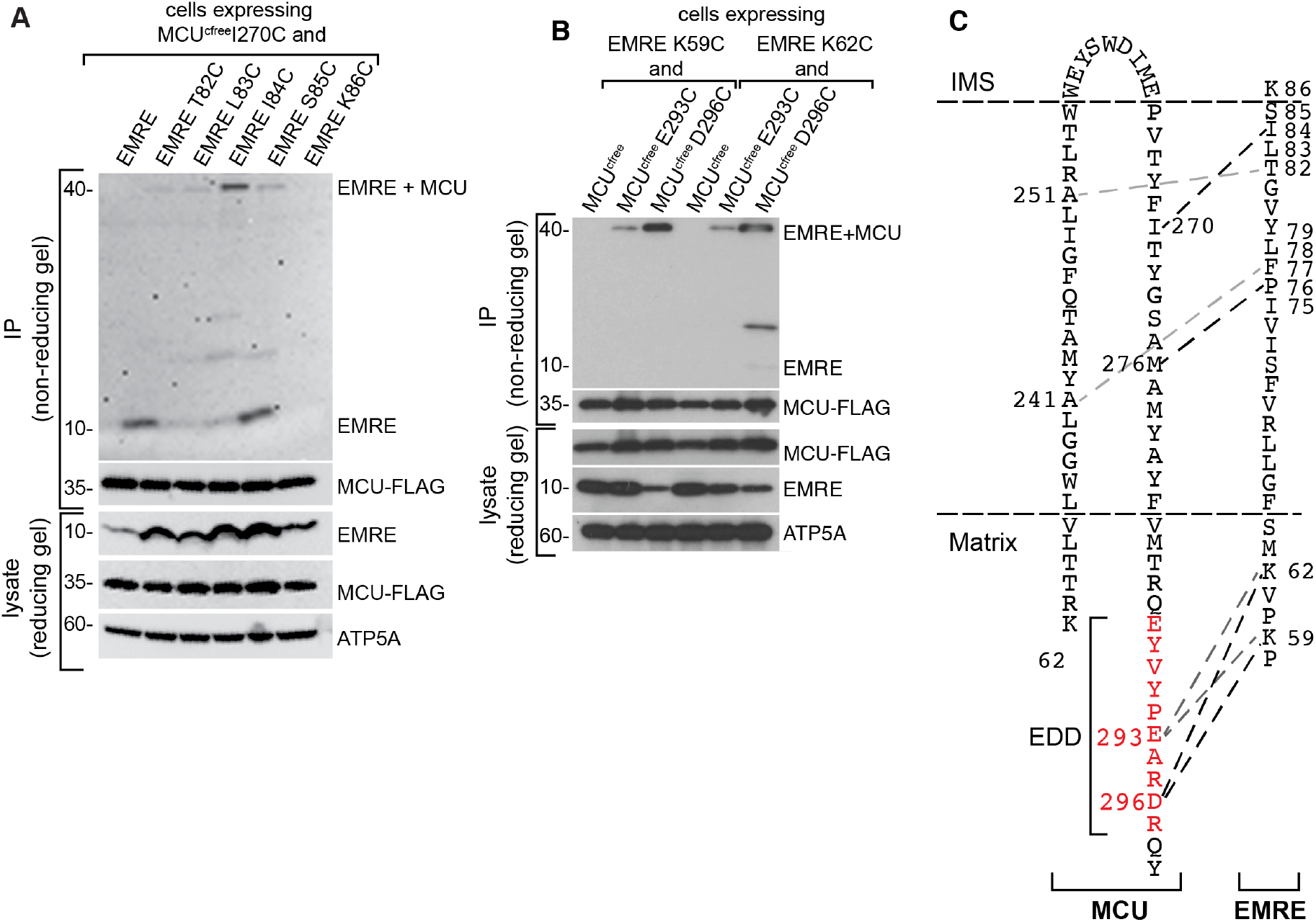
EMRE directly interacts with TM2 and EDD of MCU. (A) EMRE-HsMCU cysteine crosslinking experiment shows direct binding of MCU TM2 residue I270 to EMRE I84. HsMCU that contains only one cysteine at amino acid 270 was stably co expressed with indicated EMRE proteins in MCU KO cells. WT EMRE does not contain any cysteines and served as a control. Mitochondria were isolated from cells and cysteine-cysteine crosslinking was induced using copper phenanthroline. MCU-FLAG was immunoprecipitated and the presence of a ~40kDa crosslinked EMRE-MCU band was detected under non-reducing conditions by western blotting using EMRE antibody. Lysates were prepared in parallel under reducing conditions and were blotted to detect indicated proteins. ATP5A serves as loading control. Numbers indicate the locations of molecular weight standards. (B) EMRE-HsMCU cysteine crosslinking experiments show direct binding of MCU EDD residues E293 and D296 to EMRE K59 and K62. HsMCU proteins with one cysteine at amino acids 293 or 296 were stably co-expressed with indicated EMRE proteins in MCU KO cells. Cysteine-free MCU served as a control. Mitochondria were isolated from cells and cysteine-cysteine crosslinking was induced using copper phenanthroline. MCU-FLAG was immunoprecipitated and the presence of a ~40kDa crosslinked EMRE-MCU band was detected under non-reducing conditions by western blotting using EMRE antibody. Lysates were prepared in parallel under reducing conditions and were blotted to detect indicated proteins. ATP5A serves as a loading control. Numbers indicate the locations of molecular weight standards. (C) Schematic showing MCU and EMRE amino acids that directly interact with each other in the membrane and in the matrix. EDD is shown in red.

First, we co-expressed MCU and EMRE proteins that contain one cysteine residue each, as well as control cysteine-free MCU, in MCU KO cells and confirmed that they are functional (**Supplementary Figure 6A).** WT EMRE does not contain any cysteine residues and served as an additional control. Then, we performed copper-mediated cysteine crosslinking experiments in mitochondria isolated from these cells, immunoprecipitated MCU and determined the presence of EMRE-MCU crosslinked protein product using non-reducing gel electrophoresis followed by western blotting. Our results showed that MCU TM1 residue A241C crosslinks with EMRE F77C, but not with four other EMRE residues near F77C (Figure 3C). Similarly, MCU TM1 residue A251C specifically crosslinked to EMRE T82C (Figure 3D).

To determine if EMRE also interacts with the pore forming TM2 of MCU, we performed similar crosslinking experiments with MCU I270C. MCU KO cells expressing these MCU and EMRE cysteine crosslinked residues showed Ca^2+^ uptake, showing that cysteine substitutions did not alter protein function **(Supplementary data 6B).** MCU I270C crosslinked to EMRE I84C (Figure 4A). Collectively, these findings confirm the interaction that was observed between MCU TM1 and EMRE previously (M. F. Tsai et al., 2016), and also establish that the pore forming TM2 of MCU also interacts with EMRE.

Finally, we performed cysteine crosslinking experiments between amino acids in EDD (MCU E293 and D296) and the N-terminal domain of EMRE that faces the IMS (K59 and K62). First, we confirmed that cysteine substitutions did not alter the function of MCU or EMRE **(Supplementary Figure 6C).** Both MCU residues in EDD crosslinked to both EMRE residues (Figure 4C), suggesting that there may be flexibility in this region of the complex. Figure 4D shows MCU and EMRE amino acids that are in close contact with each other. Based on these functional and biochemical experiments, we propose that EMRE directly interacts with both TM1 and TM2 of MCU in the membrane, and also binds to EDD of MCU.

To understand the significance of EMRE-EDD interaction for uniporter function, we identified the homologous EDD region in four fungal MCU homologs whose high-resolution structures have been published and highlighted EDD in these structures (**Supplementary Figure 7)**. Surprisingly, EDD appeared flexible in these structures. Based on this observation, we posit that binding of EMRE stabilizes this otherwise flexible region and allows exit of Ca^2+^ ions from the pore.

## Discussion

Perturbation of uniporter function is associated with a number of cellular and systemic defects, ranging from altered cell cycle progression and mitochondrial dynamics to neurodegenerative disease and cardiomyopathy (Chakraborty et al., 2017; Kamer & Mootha, 2015; Musa et al., 2019). EMRE has emerged as a core component of the animal mitochondrial Ca^2+^ uniporters whose expression is under transcriptional and post-translational control (Konig et al., 2016; Munch & Harper, 2016; C. W. Tsai et al., 2017). For example, accumulation of EMRE protein in the absence of mitochondrial AAA-proteases AFG3L2 and SPG7, whose mutations are associated with spinocerebellar ataxia and hereditary spastic paraplegia, is responsible for mitochondrial Ca^2+^ overload and may contribute to neuronal loss. In addition, in a mouse model of neuromuscular disease caused by MICU1 deficiency, decreased EMRE expression over time correlated with improved health (Liu et al., 2016). These observations highlight the importance of EMRE in physiology and disease.

High-resolution structures of MCU homologs from fungal species showed a homotetrameric Ca^2+^ pore formed by the second TM domain (TM2) of MCU, and also established that MCU has a unique channel architecture among Ca^2+^ channels. Animal uniporter differs from its fungal and plant counterparts in terms of its biophysical properties, protein composition and likely in its regulation. Despite significant progress made in understanding the regulation and structure of MCU, MICU1 and MICU2, no structural data exists for EMRE, and molecular basis of EMRE function remains unknown.

Here, we exploited evolutionary divergence of mitochondrial Ca^2+^ uniporter composition to understand the function of EMRE, a metazoan specific protein. Functional experiments using chimeric proteins that consist of human HsMCU (which is EMRE-dependent) and Dictyostelium DdMCU (which operates independent of EMRE) revealed the presence of a region in MCU that we named EMRE dependence domain (EDD). We also show that EMRE makes direct contacts with the two TM domains of MCU as well as with EDD. Interestingly, the region that corresponds to EDD appears flexible in previously published high-resolution structures of MCU homologs, suggesting that binding of EMRE stabilizes this region and enables Ca^2+^ conductance. In species that do not have EMRE, it is plausible that lipids or other currently unknown proteins may fulfill the same function. It is notable that fungal MCU homologs appear to have extremely low conductance, distinct from Dictyostelium and animals, as described by Lehningner and colleagues (Carafoli & Lehninger, 1971) and subsequently by our group (Kovacs-Bogdan et al., 2014).

In our experimental system, we were unable to achieve comparable stable expression of our chimeric proteins and wild type controls and thus could not directly compare the activities of the chimeras to HsMCU or DdMCU. However, expression level of each chimeric protein was similar in EMRE KO or MCU KO cells, with the exception of Chimera 4. This enabled us to determine the EMRE-dependence of our chimeras. The measured activity of a chimeric protein is a function of its expression, channel properties and its effect on mitochondrial membrane potential and mitochondrial health. Although we did not observe perturbed membrane potential in cells that express chimeric proteins, it is possible that in our assays, some chimeric proteins exhibit reduced mitochondrial uptake because their stable expression causes mitochondrial Ca^2+^overload, alters mitochondrial Ca^2+^ storage capacity or Ca^2+^ extrusion from the mitochondria.

Our data show that EMRE directly interacts with MCU in TM1, TM2 and EDD. Importantly, comparison of chimera 7 and chimera 9 immunoprecipitation data (Figure 3A) suggests that the region defined by Chimera 10 might also be important for EMRE-MCU binding. However, Chimera 10 was not functional, and we did not pursue this chimera for protein-protein interaction experiments due to the possibility that it is not folded properly. In addition, although EDD is the smallest region that we tested that enabled EMRE-independent Ca^2+^ uptake, it is evident in our data that the presence of EMRE can increase Ca^2+^ uptake rates by Chimera 9. Thus, we note that the data presented here are consistent with additional, extended interactions between EMRE and MCU in the matrix.

What is the evolutionary significance EMRE? Curiously, based on functional data, *Dictyostelium* and fungal MCU appear to be able to adopt an open conformation when expressed (Baradaran et al., 2018; Fan et al., 2018; Kovacs-Bogdan et al., 2014; Nguyen et al., 2018; Yoo et al., 2018), whereas human MCU when expressed on its own adopts a closed conformation. The current works suggests that the EDD of MCU is responsible for maintaining the closed state, in a way that is dependent on EMRE. In addition to maintaining MCU in an open conformation, we previously showed that EMRE is also crucial for interaction with MICU1. We speculate that EMRE is responsible for transducing Ca^2+^ sensing by MICU1/2 to the EDD to gate the activity of the pore distal to the selectivity filter. If true, this model would be distinct from recent studies that have suggested that MICU1/2 directly block the uniporter at the pore opening. It is notable that the expression of EMRE has emerged as an important determinant of uniporter activity (Konig et al., 2016; C. W. Tsai et al., 2017), suggesting that this metazoan innovation may have evolved to confer an added layer of acute and chronic regulation to the uniporter. Future structural and functional studies will be required to fully decipher the mechanisms by which the uniporter is regulated across different eukaryotes.

## ACKNOWLEDGEMENTS

We thank all members of the Mootha and Sancak laboratories for critical reading of the manuscript. We also thank David M. Shechner for help with Pymol, analysis of published MCU structures and with gRNA expression advice.

## AUTHOR CONTRIBUTIONS

M.J.S.M, V.K.M and Y.S designed experiments and wrote the manuscript. A.L.M contributed to cloning of constructs, cell line maintenance, mitochondrial calcium uptake assays, crosslinking experiments. M.B., F.B. and O.G. contributed to cloning and immunoprecipitation experiments and cell line generation and maintenance.

