## Supplementary Figures for "Molecular basis of EMRE-dependence of the human mitochondrial calcium uniporter"

**Supplementary Figure 1**

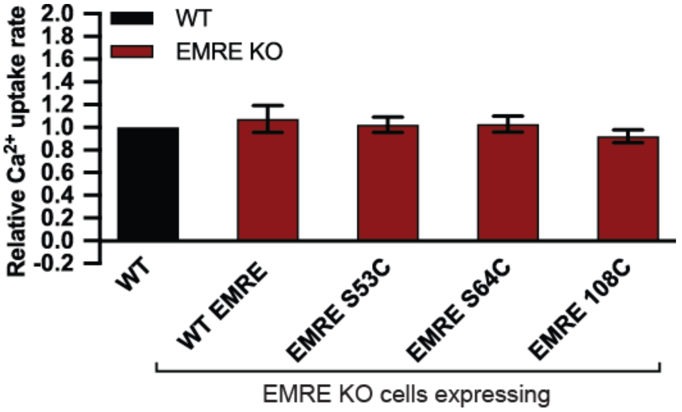


**Supplementary Figure 1: EMRE S53C, S64C and 108C are functional proteins**
Mutation of EMRE S53 and S64C to cysteine or addition of a cysteine at the C-terminus of EMRE does not impair its function. HEK293T cells expressing indicated proteins were permeabilized and mitochondiral calcium uptake was measured by monitoring extramitochondrial $Ca^{2+}$ clearance. Bar graph shows quantification of $Ca^{2+}$ uptake rates relative to WT HEK293T cells.

**Supplementary Figure 2**

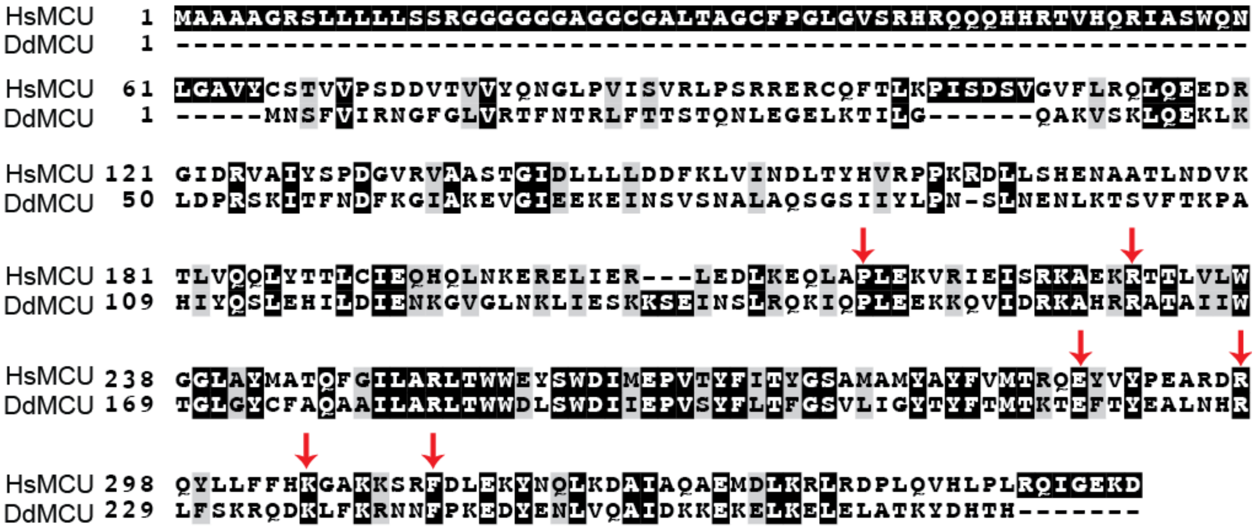


**Supplementary Figure 2: HsMCU-DdMCU chimera breakpoints**
Arrowheads point to the conserved amino acids of HsMCU and DdMCU that were chosen as break points in chimeras. Alignment of HsMCU and DdMCU was done using CLUSTALW and amino acids were color coded using Boxshade. Black boxes show identical amino acids, gray boxes show similar amino acids.

### Supplementary Figure 3

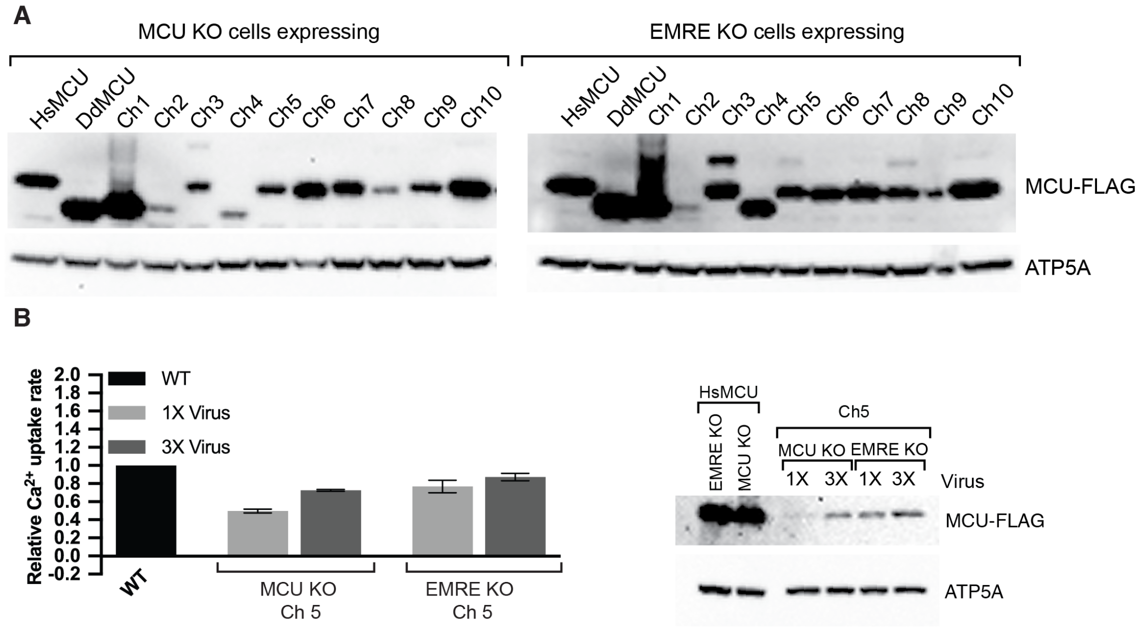


**Supplementary Figure 3: Comparison of HsMCU-DdMCU chimera protein expression and function**

(A) MCU KO or EMRE KO cells that stably express the indicated FLAG-tagged proteins were lysed, and lysates were analyzed by western blotting using anti-FLAG antibody. ATP5A serves as loading control.

(B) MCU KO and EMRE KO cells were infected with virus as described in material and methods to generate 1X infected cells after puromycin selection. The same cells were infected again with twice the amount of original virus, to produce 3X infected cells. Mitochondrial calcium uptake rates of control WT, 1X infected and 3X infected cells were measured. Bar graph shows mitochondrial $Ca^{2+}$ uptake rates relative to WT control cells. Error bars report -/+ S.D. n=3. Western blot shows expression levels of FLAG-tagged HsMCU and chimera 5. ATP5A serves as loading control.

**Supplementary Figure 4**

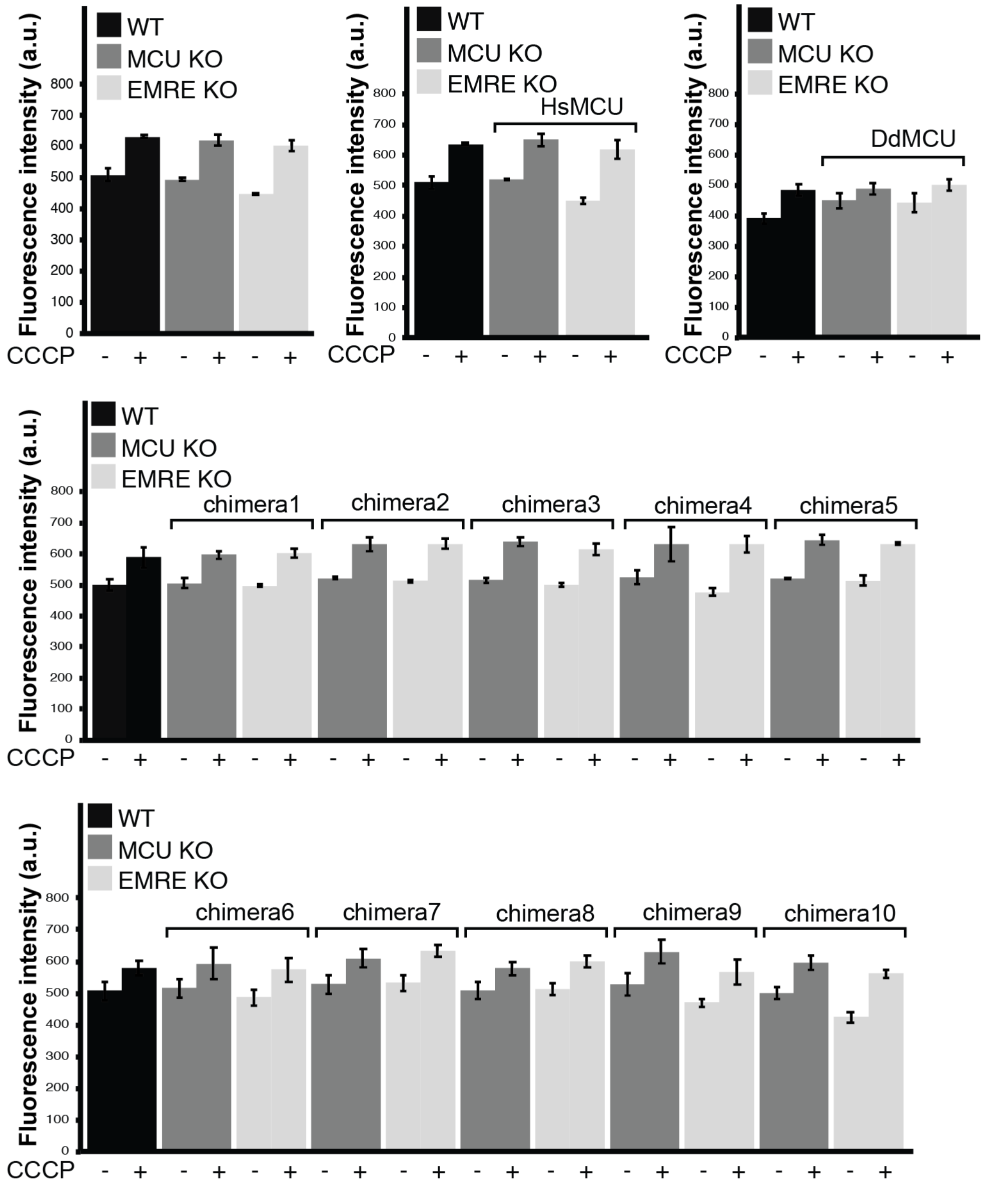


**Supplementary Figure 4: Expression of HsMCU, DdMCU or chimeras do not alter mitochondrial membrane potential**

Control WT, MCU KO or EMRE KO cells expressing the indicated proteins were permeabilized and incubated with TMRM. TMRM fluorescence was measured in the absence and presence of uncoupler CCCP. All cells showed comparable mitochondrial membrane potential. Error bars report -/+ S.D. n=3

**Supplementary Figure 5**

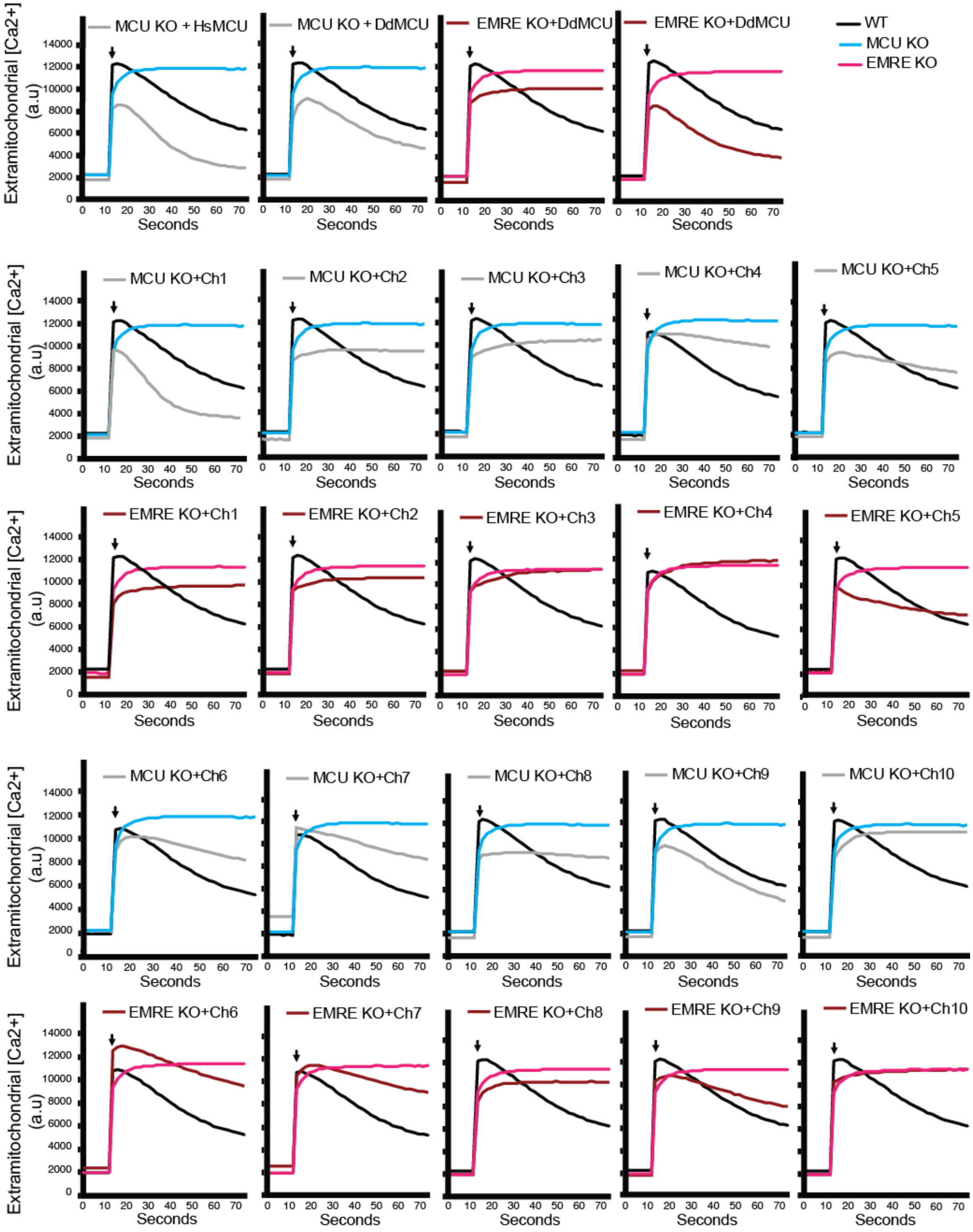


**Supplementary Figure 5: Representative mitochondrial $Ca^{2+}$ uptake traces**
WT, MCU KO or EMRE KO HEK293T cell expressing the indicated proteins were permeabilized and mitochondrial $Ca^{2+}$ uptake was measured by monitoring extramitochondrial $Ca^{2+}$ clearance. The arrows indicate addition of calcium. Each graph shows data from WT and MCU KO or EMRE KO cells, in addition to data from cell lines that express the indicated chimeras.

### Supplementary Figure 6 A,B,C

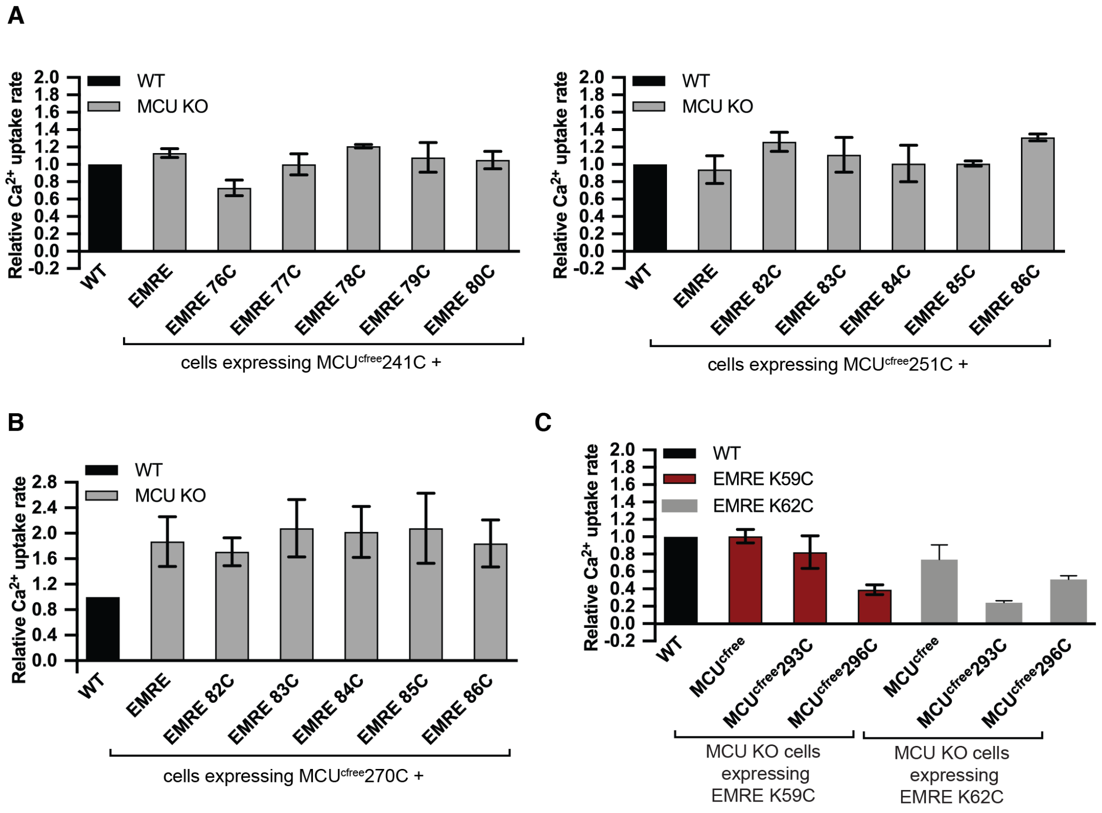


**Supplementary Figure 6: Functional characterization of MCU and EMRE proteins used for cysteine-crosslinking experiments**

MCU KO HEK293T cells expressing indicated proteins were permeabilized and mitochondrial $Ca^{2+}$ uptake was measured by monitoring extramitochondrial $Ca^{2+}$ clearance. Bar graphs show $Ca^{2+}$ uptake rates relative to control WT cells. Error bars report -/+ S.D, n=3-4

(A) MCU-EMRE crosslinking in TM1 of MCU

(B) MCU-EMRE crosslinking in TM2 or MCU

(C) MCU-EMRE crosslinking in EDD of MCU

**Supplementary Figure 7**

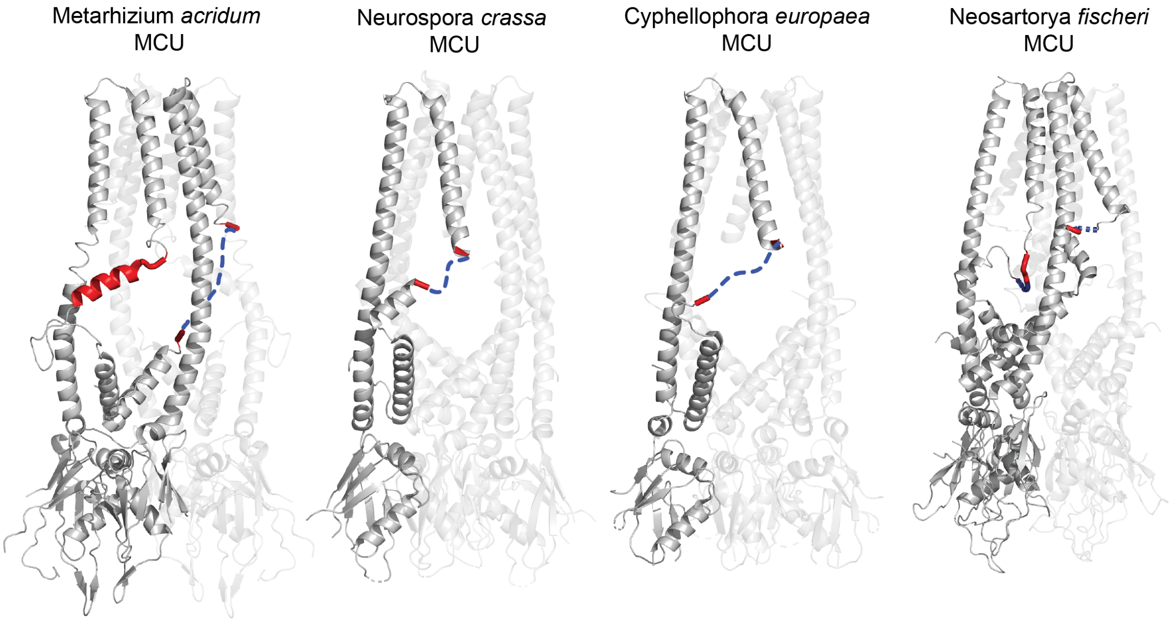


**Supplementary Figure 7: The region homologous to EDD appears flexible in fungi**
High-resolution structures of MCU tetramers from four different fungal species are shown. The region that is homologous to EDD or amino acids that surround the EDD in these structures are shown in red. Dotted blue lines show amino acids that are omitted in these structures, all of which overlap with EDD. In the *M. acridum* MCU structure, one MCU chain shows alpha helical EDD, whereas the same region in the neighboring chain is not structured. PDB IDs of these structures are as follows: *M. acridum* (6C5W); *N. crassa* (5KUJ); *N. fischeri* (6D7W); *C. europaea* (6DNF)
